# Histone dynamics within the nucleosome play a critical role in SNF2h-mediated nucleosome sliding

**DOI:** 10.1101/2020.03.14.992073

**Authors:** Nathan Gamarra, Geeta J. Narlikar

**Affiliations:** Department of Biochemistry and Biophysics, University of California, San Francisco, San Francisco, United States; TETRAD Graduate Program, University of California, San Francisco, San Francisco, United States

## Abstract

Elucidating the mechanisms by which ATP-dependent chromatin remodeling enzymes disrupt nucleosome structure is essential to understanding how chromatin states are established and maintained. A key finding informing remodeler mechanism is the observation that the dynamics of protein residues buried within the histone core of the nucleosome are used by specific remodelers to mobilize the nucleosome^1^. Recently, a study obtaining cryo-electron microscopy (cryo-EM) structures of ISWI-family remodeler-nucleosome complexes failed to observe stable conformational rearrangements in the histone octamer^2^. The authors of this study also failed to replicate the earlier finding that site-specifically restraining histone dynamics inhibits nucleosome sliding by ISWI-family remodelers^1,2^. In contrast, a recent cryo-EM structure detected asymmetric histone dynamics in an ISWI-nucleosome complex^3^. Here, using two different protocols, we replicate the original finding in Sinha et al.^1^ that dynamics within the histone core are important for nucleosome sliding by the human ISWI remodeler, SNF2h. These results firmly establish histone dynamics as an essential feature of ISWI-mediated nucleosome sliding and highlight the care required in designing and performing biochemical experiments investigating nucleosome dynamics using disulfide linkages.

## Results and Discussion

In previous work, dynamics in the histone core were demonstrated to be functionally important for nucleosome sliding by SNF2h using site-specific disulfide crosslinks engineered in otherwise cysteine-free histones^1^. These crosslinks inhibited nucleosome sliding by SNF2h, suggesting that inhibition of sliding was due to restraining histone octamer dynamics. To minimize any potential damage to the nucleosome that could influence the experimental results, oxidation reactions for generating the disulfide bond were carried out in the absence of metal catalysts, which can generate free radicals that may create non-specific damage^4^. Since the publication of this work, additional studies investigating both ISWI-mediated^5^ and spontaneous^6^ nucleosome sliding using disulfide crosslinking in the absence of metal catalysts have reinforced the view that histone conformational dynamics play an important role in nucleosome sliding.

In contrast, Yan et al.^2^ reported that restraining histone dynamics using single H3-H4 crosslinks (H3L82C-H4V81C called sCX2) from Sinha et al.^1^ did not specifically impact nucleosome sliding by SNF2h. Yan et al. first generated sCX2 crosslinks in the background of WT *Xenopus laevis* octamer, which bears an endogenous cysteine (H3C110) and found that sliding was unaffected. However, the presence of an additional reactive cysteine, which has previously been shown to readily form disulfides under oxidizing conditions^7^, creates the possibility for multiple types of crosslinked species, complicating the interpretation of these results. Yan et al. also created sCX2 crosslinks in an H3C110A octamer and, as in Sinha et al., found that SNF2h sliding activity was impaired, but to a much lesser extent than observed previously. Further, when the disulfide bond was apparently reduced by adding 100mM DTT, sliding activity was not restored. Based on these results, Yan et al. suggested, “…that the loss of activity did not result from the disulfide bond but probably from non-specific oxidation damage to the nucleosome, which was prone to precipitate out of the solution under the condition of extensive oxidation. Prolonged treatment might have resulted in damage to the nucleosome and loss of the activity.” However, the authors did not directly test for this possibility as they did not report any remodeling experiments with nucleosomes lacking the cysteine residues that had been subjected to the same oxidation protocol.

We therefore set out to directly test the possibility raised by Yan et al.^2^ of nonspecific oxidative damage to the histone octamer and to determine whether any differences in the protocols used to generate crosslinked nucleosomes could explain the discrepancies between the two studies. First, we assembled nucleosomes with *Xenopus laevis* H3C110A octamer that had been oxidized with CuPhe (Supplemental Figure 1A). No apparent precipitates were observed during the preparation of oxidized octamer. Oxidized octamer also readily assembled into canonical nucleosomes, suggesting no gross defects. These oxidized nucleosomes were then purified via glycerol gradient ultracentrifugation and were remodeled with SNF2h. No defect was seen in the remodeling of nucleosomes assembled from oxidized H3C110A octamers when compared to nucleosomes assembled from untreated H3C110A octamers (Table 1, Figure 1A). This result indicates that non-specific oxidative damage to the octamer cannot explain the remodeling defects.

**Figure 1.**
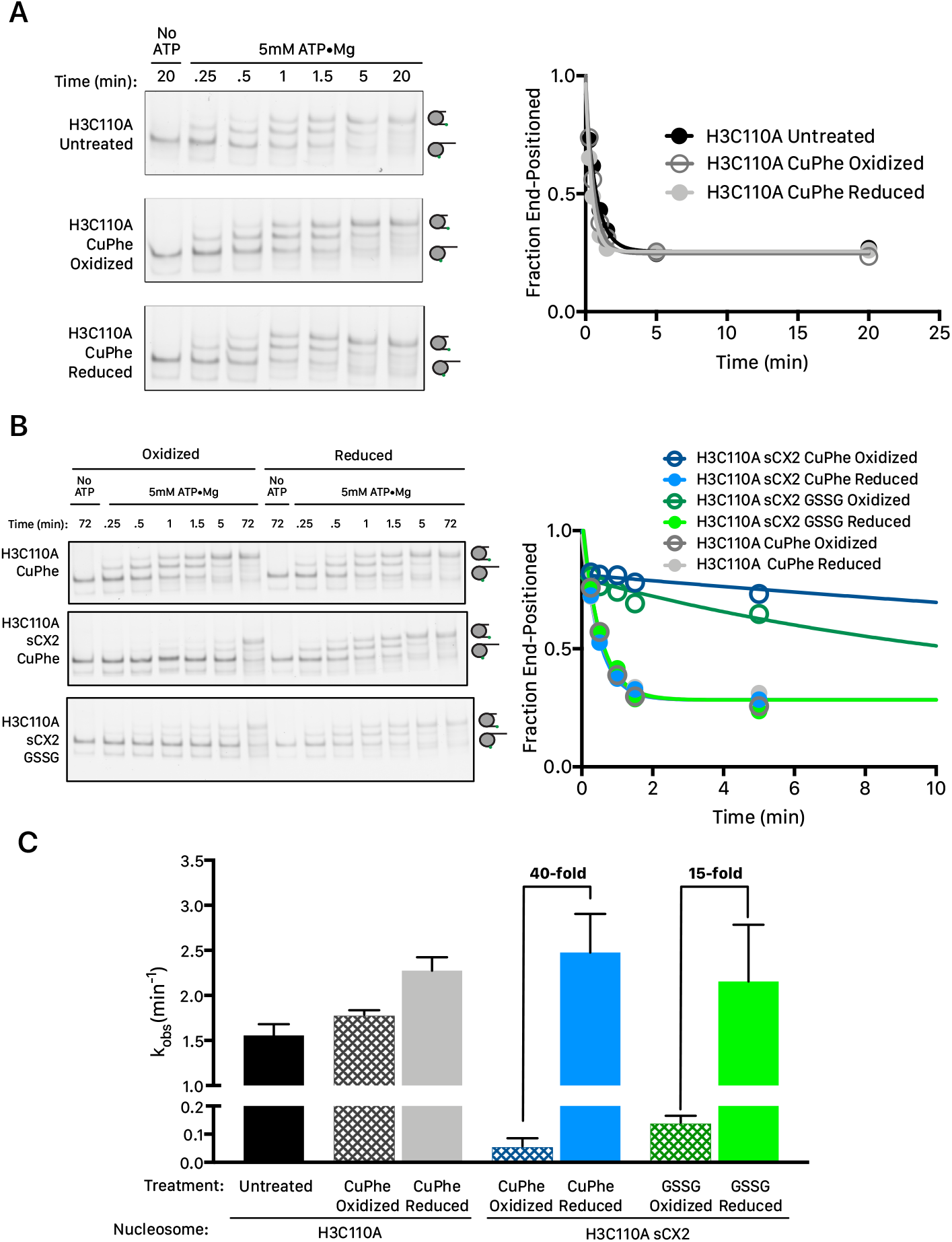
Site-specific cysteine crosslinking, and not general oxidative damage, inhibits remodeling regardless of oxidation method. (A) Treatment of H3C110A histone octamer with CuPhe or GSSG does not affect nucleosome sliding by SNF2h. Left. Native gel sliding assay with saturating concentrations of SNF2h (1μM) with or without saturating ATP and 15nM cy3-labeled nucleosomes using the indicated histone octamer. Time points were quenched with excess ADP and plasmid DNA and resolved on a 6% (29:1 bis) acrylamide gel. Higher migrating species are more centrally-positioned nucleosomes. Right. Quantification of the gel data plotted as the fraction of end-positioned nucleosomes over time. (B) Native gel remodeling assay as in A with the plot zoomed in to the first 10 minutes of the reaction to better evaluate fits. Nucleosomes assembled with oxidized H3C110A sCX2 octamers are slid substantially slower than nucleosomes assembled with oxidized H3C110A octamers. This effect can be completely reversed by treatment of the oxidized H3C110A sCX2 octamer with reducing agent prior to nucleosome assembly. (C) Mean observed rate constants (k_obs_) from 3 independent experiments. Error bars reflect the standard error of the mean (SEM).

**Table 1.** k_obs_ for nucleosome sliding of the given substrate at different concentrations of SNF2h under buffer conditions from Sinha et al.^1^

| Octamer used to assemble nucleosome | **k<sub>obs</sub>, number of replicates** |  |
| --- | --- | --- |
|  | **[SNF2h]** |  |
|  | **1μM** | **50nM** |
| **H3C110A Untreated** | 1.3±0.2 min<sup>-1</sup>, n=6 | 0.03±0.01 min<sup>-1</sup>, n=3 |
| **H3C110A CuPhe Oxidized** | 1.8±0.1 min<sup>-1</sup>, n=3 | Not Determined |
| **H3C110A CuPhe Oxidized and then reduced** | 2.3±0.1 min<sup>-1</sup>, n=3 | Not Determined |
| **H3C110A/L82C CuPhe Oxidized** | 0.63±0.05 min<sup>-1</sup>, n=3 | Not Determined |
| **H3C110A/H4L81C CuPhe Oxidized** | 0.54±0.04 min<sup>-1</sup>, n=3 | Not Determined |
| **H3C110A sCX2 CuPhe Oxidized** | 0.05±0.01 min<sup>-1</sup>, n=6 | Undetectable, n=3 |
| **H3C110A sCX2 CuPhe Oxidized and then reduced** | 2.5±0.4 min<sup>-1</sup>, n=3 | 0.03±0.02 min<sup>-1</sup>, n=3 |
| **H3C110A sCX2 GSSG Oxidized** | 0.14±0.02 min<sup>-1</sup>, n=3 | Not Determined |
| **H3C110A sCX2 GSSG Oxidized and then reduced** | 2.2±0.6 min<sup>-1</sup>, n=3 | Not Determined |

We then wondered whether the oxidation method may impact the crosslinking results only in the context of H3C110A sCX2 octamer. To test this possibility, we generated H3C110A sCX2 crosslinked nucleosomes using CuPhe or using oxidized glutathione (GSSG), a gentler oxidizing agent that does not generate free radicals and has been used previously to investigate nucleosome dynamics^8^. Oxidation with CuPhe went to near-completion as assessed by a non-reducing SDS-PAGE gel, while GSSG oxidation was less efficient (Supplemental Figure 1A). A portion of each H3C110A sCX2 crosslinked octamer was then reduced by dialysis into high salt buffer containing 100mM DTT (Supplemental figure 1B). H3C110A sCX2 octamers also showed no visible precipitates upon oxidation or reduction and, like cysteine-free octamers, readily assembled into nucleosome core particles that were purified by glycerol gradient ultracentrifugation. H3C110A sCX2 nucleosomes assembled with CuPhe oxidized octamer were slowed in nucleosome sliding ~40-fold compared to reduced controls under saturating SNF2h conditions, while GSSG octamer slowed sliding ~15-fold (Figures 1B and C). The difference in quantitative effects between the two oxidation methods can be attributed to the different crosslinking efficiencies in each sample and cannot be explained by nucleosome disassembly (Supplemental Figures 2 and 3). Reduction of either type of oxidized octamer sample fully restored sliding activity in nucleosomes assembled from the respective octamers (Figures 1B and C). To further test whether the oxidation-dependent slowing of sCX2 nucleosomes is due to the formation of a disulfide bond, we also tested remodeling on nucleosomes assembled using oxidized H3C110A octamers containing single cysteines from the sCX2 cysteine pair that were purified by ultracentrifugation (Supplemental Figure 4A). In this side-by-side experiment, while CuPhe oxidized H3C110A sCX2 nucleosomes remodeled ~30-fold slower than untreated H3C110A nucleosomes, single cysteine-containing nucleosomes remodeled only ~2-fold slower. Oxidation-dependent remodeling defects were also observed with sub-saturating concentrations of enzyme (Supplemental Figure 4B). Additionally, the remodeling conditions of Yan et al. ^2^ differ from Sinha et al. (see methods). Under their remodeling conditions we found that remodeling reactions were faster overall (Supplemental Figure 4C). However, even under these conditions the disulfide bond causes a >6-fold reduction in reaction rate (Supplemental Figure 4C). Because the remodeling of uncrosslinked nucleosomes is faster under these conditions, the first time point we are able to capture is already ~60 % remodeled. As a result, the rate constant reported here is likely to be an underestimate of the actual rate constant and the 6-fold defect should be considered a lower limit. These results indicate that the sliding inhibition observed with nucleosomes assembled using the oxidized H3C110A sCX2 octamer is explained by the disulfide mediated restraint placed on histone dynamics.

The results that we describe above, (i) rule out the model put forth by Yan et al. of non-specific octamer oxidation causing inhibition of sliding; (ii) reproduce the large inhibitory effects seen by Sinha et al. when the H3L82C-H4V81C disulfide bond is formed in an H3C110A nucleosome; and (iii) show that the large defect is completely reversible when the oxidized octamer is reduced prior to nucleosome assembly. These results are summarized in Table 1.

Yan et al. carried out their test for reversibility in a different manner than Sinha et al. While they carried out the forward reaction, namely oxidation in the context of an octamer (as Sinha et al. did), they carried out the reverse reaction (reduction) in the context of a nucleosome. In our experience, reducing buried disulfides in the context of a fully assembled nucleosome has been very difficult and we reason this is because the disulfide crosslink becomes significantly less solvent accessible in a nucleosome compared to an octamer. However, to complete our analysis of the discrepancy between Sinha et al., and Yan et al., we decided to test the method used by Yan et al. In their method for reducing the disulfide bonds, Yan et al. treated the crosslinked nucleosomes with 100mM DTT at 37°C for 1hr before initiating remodeling reactions. When we reduced crosslinked H3C110A sCX2 nucleosomes as in Yan et al., we found that sliding activity was not restored, consistent with their report (Supplemental Figure 5A). Further, Yan et al. showed that these conditions were sufficient to completely break the disulfide bond when assayed by nonreducing SDS-PAGE. However, we reasoned that if the sample was not buffer exchanged to remove excess DTT prior to denaturation in SDS-PAGE loading buffer, once-buried disulfides could become exposed upon denaturation and rapidly reduced by residual reducing agent present in the sample. This would cause an overestimate of the degree of reduction on the SDS-PAGE gel. To test for this possibility, we reduced H3C110A sCX2 crosslinked nucleosomes with 100mM DTT for 1hr at 37°C as in Supplemental Figure 5A and either directly added the sample to SDS loading buffer or quenched the residual DTT with 5-fold excess N-ethyl maleimide before SDS treatment. While the unquenched sample appears to have completely reversed the disulfide, quenching the sample after treatment appears completely oxidized (Supplemental Figure 5C). This cannot be explained by reoxidation of the disulfide bond upon quenching of reducing agent as N-ethyl maleimide would also react with cysteine thols, blocking new disulfide bond formation. In addition, oxidation of the cysteines within octamers in the absence of catalysts such as Cu-Phe or GSSG is extremely slow (≥48 hours, see materials and methods in Sinha et al.^1^). As a result, it is very likely that the apparent inability to reverse defects associated with crosslinking by Yan et al. may have been due to an inability to completely reverse crosslinking in fully assembled nucleosomes, and that the reduction they observe occurs while processing the samples for analysis by SDS-PAGE.

Importantly, our results do not invalidate the other conclusions presented in Yan et al. nor do they cast any doubt on the quality and accuracy of the structural data presented. The structures presented by Yan et al. represent only the fraction of states that can be reconstructed to high resolution. Lowly-populated or highly dynamic states would likely be lost during the averaging required for high resolution analysis. Consistent with this possibility, a recent cryo-EM study captured multiple stable ISWI-nucleosome states^3^. While the highest resolution structures showed no clear evidence of conformational dynamics, lower resolution structures showed local reduction in the cryo-EM density. This local reduction overlaps both with dynamic regions captured by NMR in Sinha et al., and with the crosslinked residues tested here and in Sinha et al^1^. These results are consistent with SNF2h promoting the formation of an ensemble of highly dynamic conformations that cannot be directly visualized by cryo-EM averaging. Thus, the absence of a stable deformed histone octamer in the cryo-EM density reported by Yan et al. is easily compatible with the finding that histone deformation is important for nucleosome sliding.

In summary, we replicate the findings of Sinha et al.^1^ that restraining histone dynamics with disulfide crosslinks substantially interferes with SNF2h-mediated nucleosome sliding. We further show that this finding is robust to the method of crosslinking and cannot be explained by non-specific oxidative damage to the nucleosome. The inability to replicate this finding by Yan et al.^2^ is likely in part due to incomplete reduction of the disulfide bond in the context of the nucleosome.

Overall, our results are consistent with an important role for histone octamer plasticity during SNF2h-mediated nucleosome sliding. Further, these results underscore the importance of performing both oxidation and reduction reactions in the context of the histone octamer when attempting to generate or reverse crosslinks to investigate the role of histone dynamics.

## Acknowledgements

We thank Kalyan Sinha for help generating the disulfide crosslinked nucleosome and Julia Tretyakova for preparing histones. We also thank Serena Sanulli, John Gross, Joseph Lobel, and Yifan Cheng for helpful feedback in preparing this manuscript and all members of the Narlikar lab for helpful discussions. This work was supported by a grant from the NIH to GJN (R35GM127020) and an NSF predoctoral and UCSF discovery fellowship to NG.

## Supplemental Methods

SNF2h was purified from E. coli as described previously^3^. Recombinant *Xenopus laevis* histones were expressed and purified from E. coli as previously described^9^. Cysteine mutant histones were purified in the presence of excess DTT. Mutant histones were also purified using only the gel-filtration step because cyanate formed in the urea buffers required for ion exchange reacts with the cysteine thiols^1^. Cy3-labeled nucleosome DNA was generated by PCR with HPLC-purified, labeled primers (IDT, Coralville, IA) and Phusion DNA polymerase using the strong, synthetic 601 nucleosome positioning sequence^10^. Nucleosome DNA was purified by native PAGE.

Histone octamer refolding was performed as described previously^3^. H3C110A sCX2 octamer was refolded in the presence of 10mM DTT to prevent disulfide formation. Refolded histone octamer and H2A/H2B dimer was purified by size exclusion chromatography with 10mM DTT. For oxidation, histone octamers and dimers were diluted to 5μM and ~200μL of sample was dialyzed twice into 1L high salt buffer without reducing agent (10mM HEPES pH 8.5, 2M NaCl, 1mM EDTA) each time overnight at 4°C.

For the CuPhe oxidation protocol, histone octamer and dimer was dialyzed overnight into 10mM HEPES pH 7.25, 2M NaCl. Histones were treated with 25μM Cu(II)SO4 and 100μM o-phenanthroline for 75 min at room temperature in the dark before being quenched with 10mM EDTA. Stocks of Cu(II)SO4 and o-phenanthroline used for oxidation were freshly dissolved in 10 mM HEPES pH 7.25, 2 M NaCl on the day of reaction. For the glutathione oxidation, histones were mixed 4:1 by volume histone:glutathione buffer (250mM Tris pH 9.0, 2M NaCl, 7.5mM Oxidized Glutathione, 2.5mM Reduced Glutathione, 5mM Benzamidine, 0.5mM Leupeptin) at room temperature in the dark for 4 nights. To check the progress of oxidation, aliquots of each reaction were quenched with 50mM iodoacetamide and run on a non-reducing SDS-PAGE gel. Disappearance of the H3 and H4 bands and appearance of a higher molecular weight species was used to determine the progress of oxidation. Quantification of crosslinking was done by measuring the intensity of the H3-H4 crosslinked band and normalizing it to the intensity of the H2A/H2B band. After oxidation, histones were dialyzed into 1L of 10mM HEPES pH 8.5, 2M NaCl, 1mM EDTA overnight at 4°C.

To reduce oxidized histones, samples were dialyzed overnight into 10mM HEPES pH 8.5, 2M NaCl, 1mM EDTA, 100mM DTT at 4°C. Reduced samples were checked using nonreducing SDS-PAGE. Loss of the high molecular weight species and reappearance of the H3 and H4 bands indicated complete reduction.

Nucleosomes were assembled using salt gradient dialysis and purified by glycerol gradient centrifugation as described previously^11^. Oxidized nucleosomes were assembled using buffers without reducing agent while reduced nucleosomes were assembled using buffers with 3mM TCEP.

All remodeling reactions were performed under single turnover conditions (enzyme in excess of nucleosomes) with saturating concentrations (1μM) or subsaturating (50nM) concentrations of SNF2h at 20°C with 15 nM nucleosomes, 12.5 mM HEPES pH 7.5, 2 mM Tris pH 7.5, 70 mM KCl, 5 mM ATP•MgCl2, 3 mM MgCl2, 0.02% NP40, and ~3%(v/v) glycerol. To assess the effects of the different buffer and temperature conditions used by Yan et al.^2^, we also measured remodeling using 50 nM SNF2h and 15 nM nucleosomes at 37°C with 20mM HEPES pH 7.5, 50mM KCl, 2 mM ATP•MgCl2, 3 mM MgCl2, 0.1mg/mL BSA, and 5%(v/v) glycerol. Reactions were started with addition of nucleosomes and time points were quenched with excess ADP and plasmid DNA. Time points were then resolved by native PAGE (6% acrylamide, 0.5XTBE) and scanned on a Typhoon variable mode imager (GE Life Sciences, Pittsburgh, PA) by scanning for fluorescent labels. Gels were then quantified by densitometry using ImageJ. The fraction of nucleosomes end-positioned (i.e. unremodeled) at a given time point was determined by the ratio of fast-migrating nucleosomes to the total nucleosome intensity. This was fit to a single exponential decay using Prism 6 (GraphPad, La Jolla, CA).

### Data availability

All relevant data are included in the paper and its supplementary information files. In addition, source data for the results presented are available from the corresponding author upon reasonable request.

**Supplemental Figure 1.**
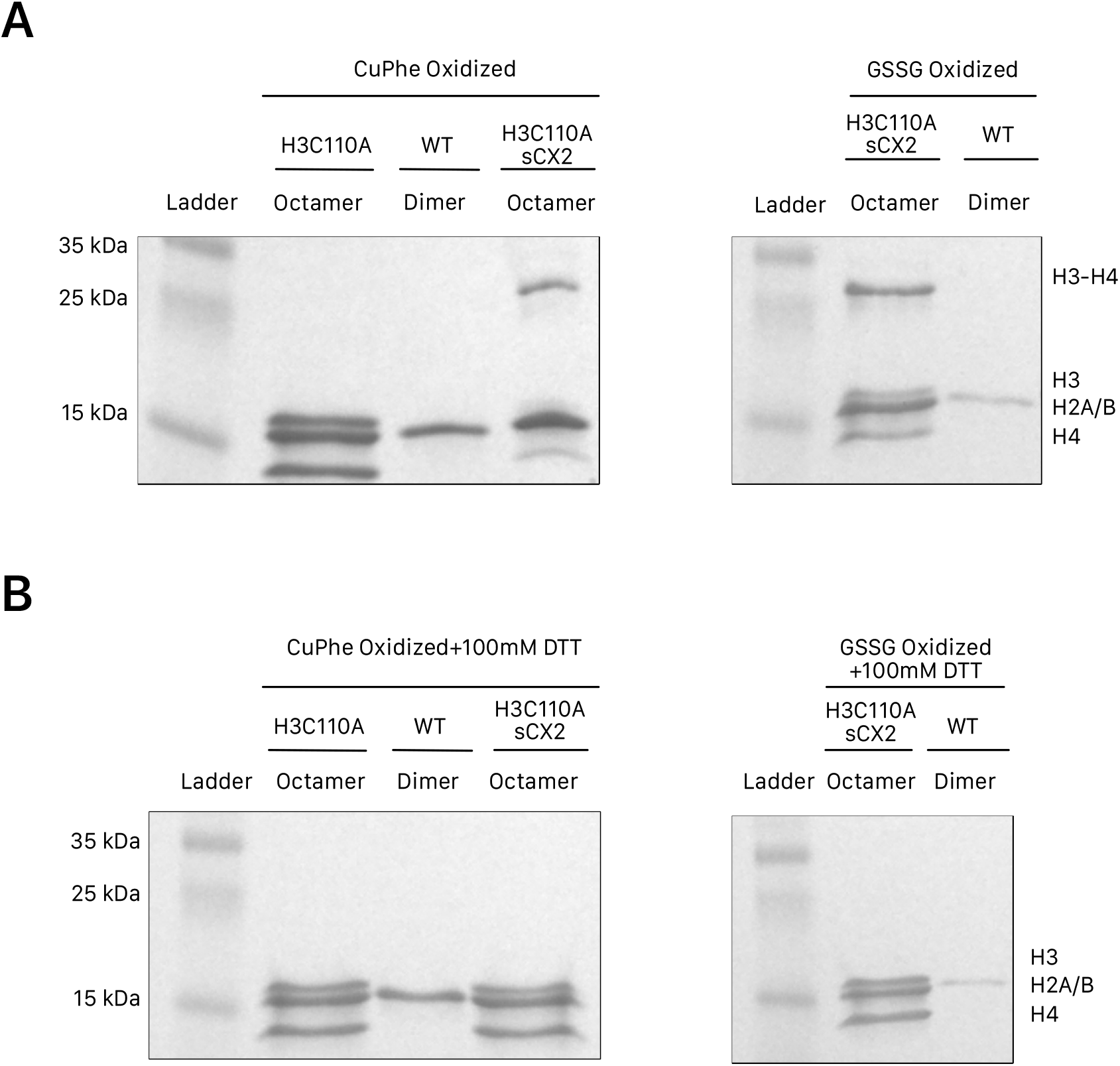
Preparation of crosslinked and reduced histone octamers. SDS-PAGE gels of *Xenopus laevis* of histones used to prepare nucleosomes in this study. (A) H3C110A histone octamer and H3C110A sCX2 (H3 L82C, H4 V81C) histone octamer oxidized using copper phenanthroline (CuPhe) or oxidized glutathione (GSSG). (B) The same samples used in (A) treated with 100mM DTT in order to reduce the disulfide bond.

**Supplemental Figure 2.**
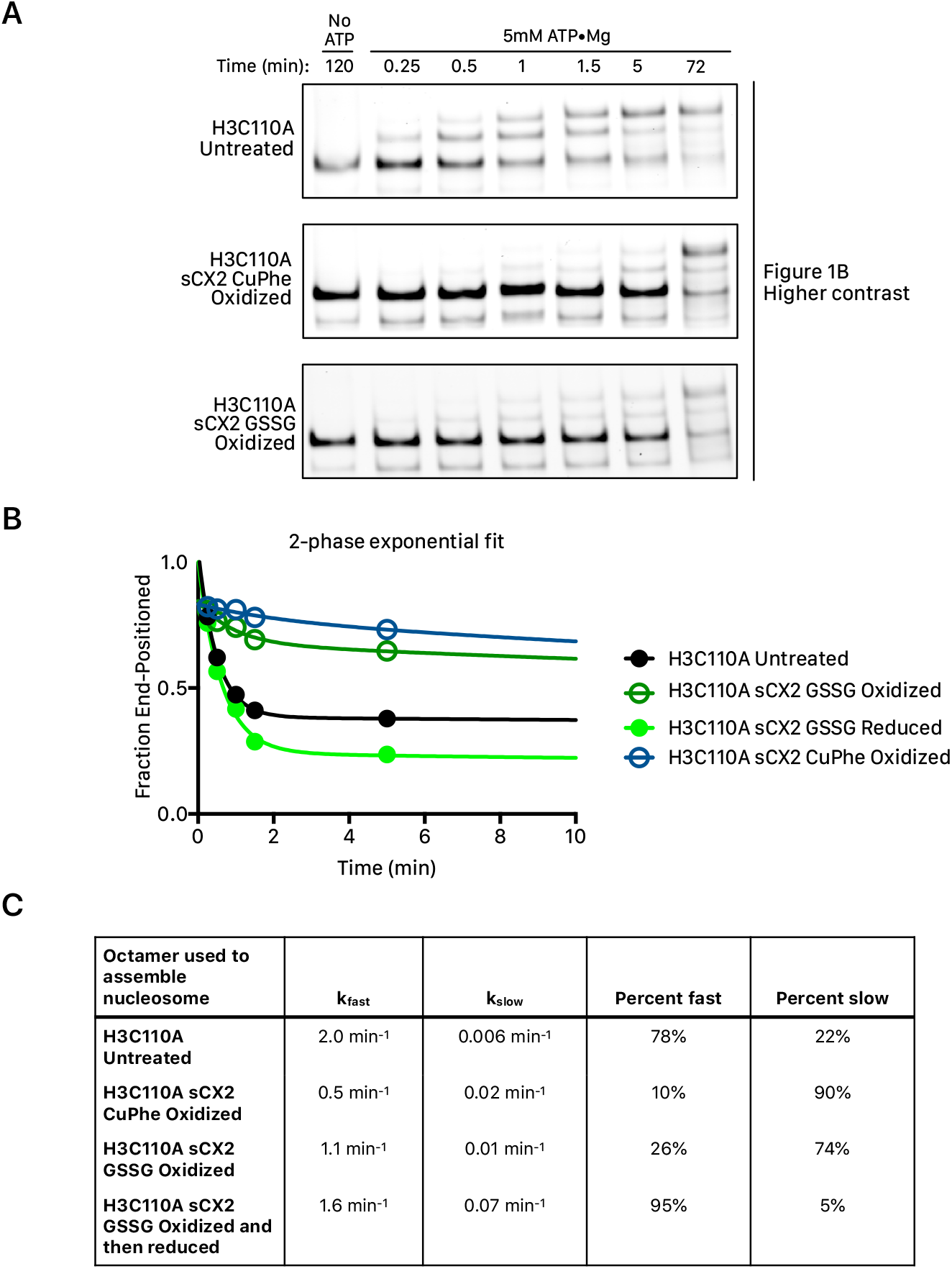
Variation in remodeling rate between oxidation methods can be explained by crosslinking efficiency. (A) Native gel remodeling assay from Figure 1B shown at higher contrast to better see the early phase of the remodeling reaction. H3C110A sCX2 GSSG oxidized nucleosomes, which have a lower crosslinking efficiency, show a population of rapidly remodeling nucleosomes. (B) Quantification of assay in A. fit to a two-phase exponential decay. (C) Best-fit parameters of the two-phase fit of the data in B. The rate constants for the fast and slow phase are similar between GSSG and CuPhe oxidized H3C110A sCX2 nucleosomes but differ in the fraction in the fast and slow phase.

**Supplemental Figure 3.**
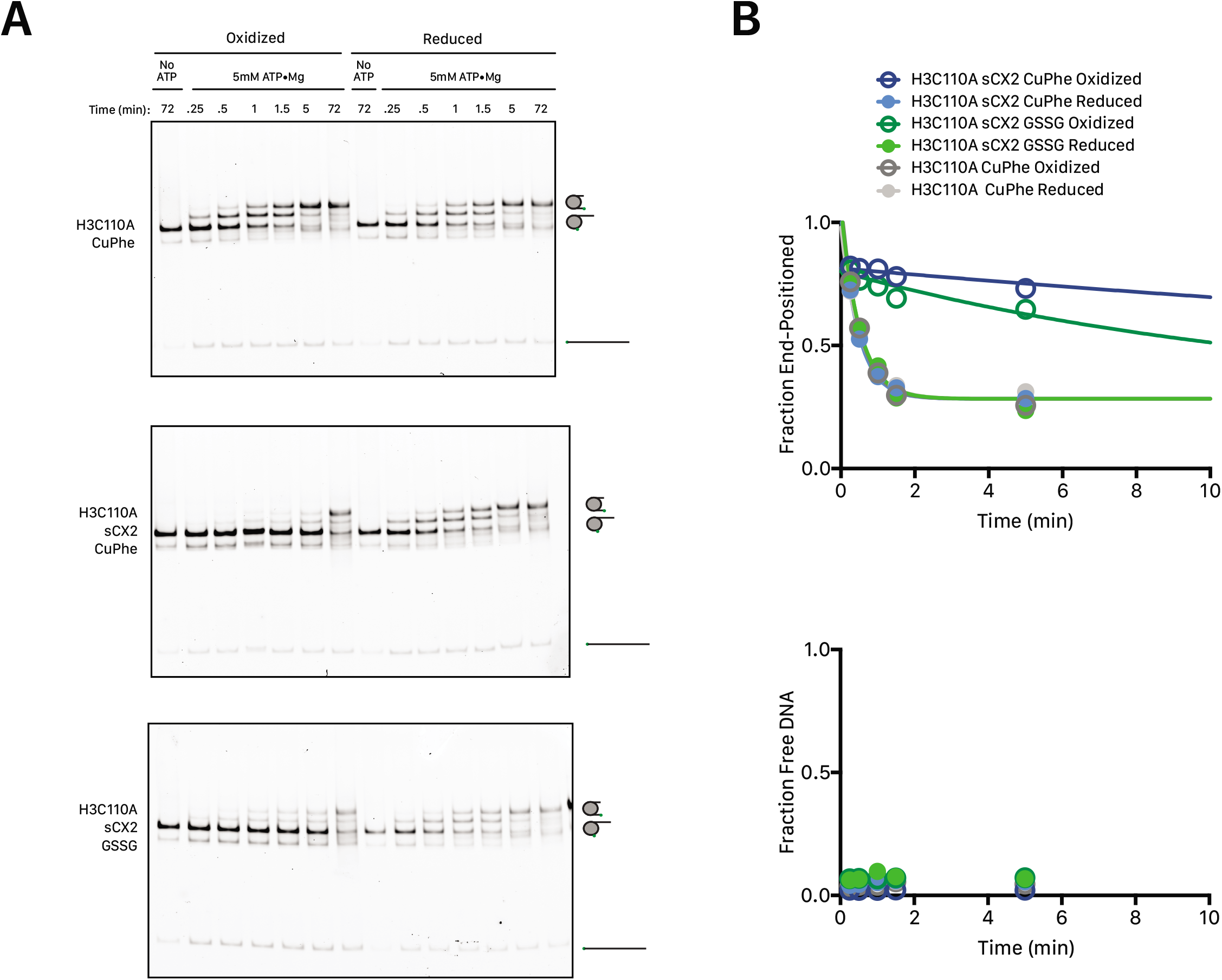
Nucleosome disassembly does not contribute to nucleosome remodeling by SNF2h. (A) Native gel remodeling data from Figure 1B of the main text presented again, but with a bigger part of the gel included to show the free DNA band on the native gel. (B) Top. Quantification of fraction end positioned nucleosomes of the gels in A and zoomed to show the first 10 minutes of the reaction as in Figure 1B. Bottom. Quantification of the fraction free DNA over time (quantified as the intensity of the free DNA band over all bands).

**Supplemental Figure 4.**
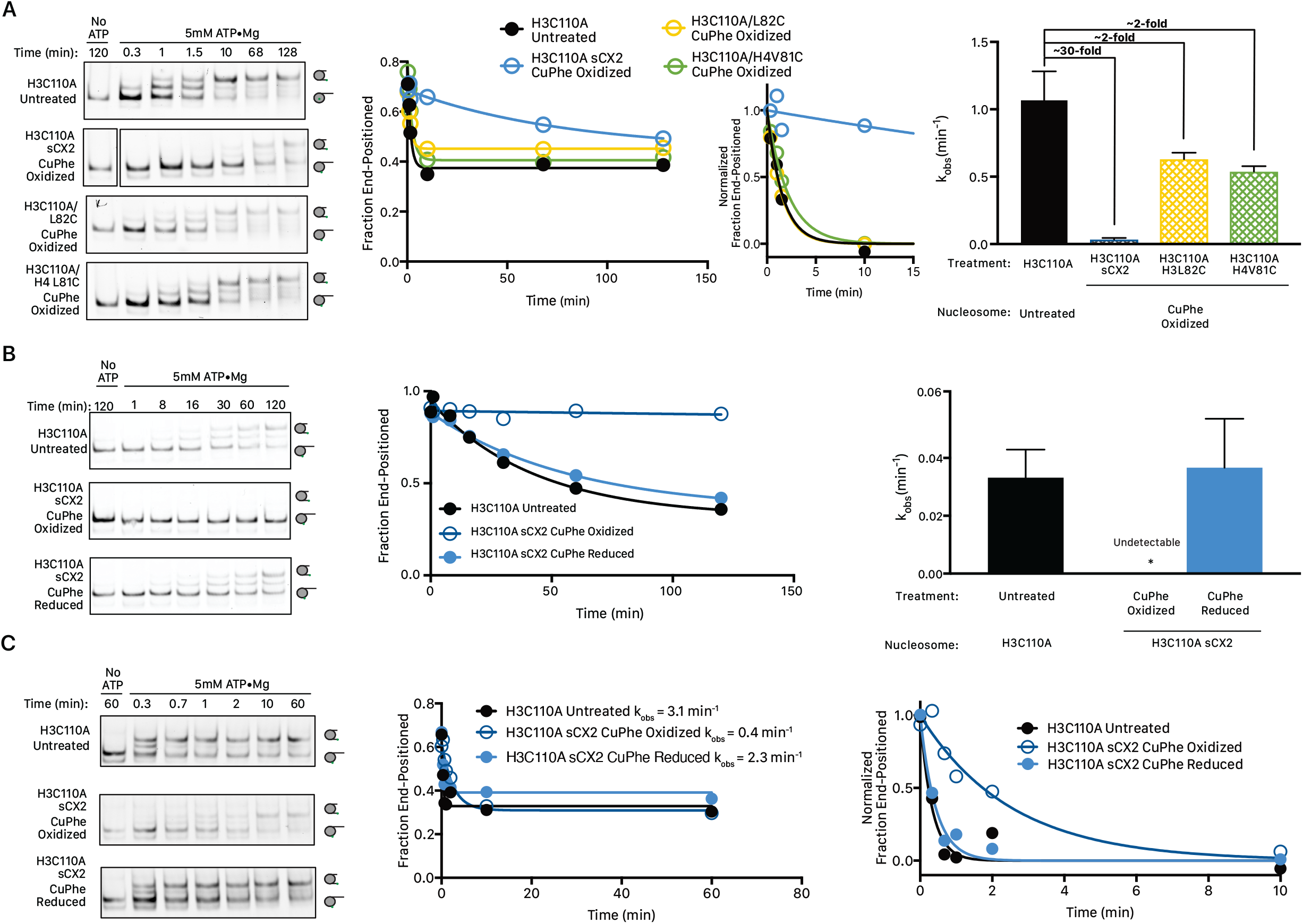
Remodeling of oxidized nucleosomes is slowed specifically due to disulfide bond formation and is robust to remodeling conditions. (A) Left. Native gel remodeling assay with saturating SNF2h (1μM), saturating ATP, and 15nM cy3-nucleosomes as in Figure 1. Nucleosomes assembled from oxidized H3C110A sCX2 octamers are remodeled substantially slower than nucleosomes assembled from untreated H3C110A octamers while oxidized octamers containing only a single cysteine are modestly slowed. Middle. Quantification of the gels on the left including a plot of all time points; and for ease of comparison a plot of the first 15 minutes of the reaction normalized to the best-fit parameters for Y_0_ and plateau. Right. Mean observed rate constants (k_obs_) from 3 independent experiments. Error bars reflect the standard error of the mean (SEM). (B) Left. Native gel remodeling assay with sub-saturating SNF2h (50nM), saturating ATP, and 15nM cy3-nucleosomes as in Figure 1. Nucleosomes assembled from oxidized H3C110A sCX2 octamers are remodeled substantially slower than nucleosomes assembled from untreated H3C110A octamers. Nucleosomes assembled from H3C110A sCX2 octamers that were first oxidized and then reduced are remodeled as fast as nucleosomes assembled from untreated H3C110A octamers. Middle. Quantification of the gels on the left. Right. Mean and SEM of the observed rate constants (k_obs_) from 3 independent experiments. The asterisk denotes that the rate constant for the oxidized reaction condition was too slow to reliably quantify with the time points taken. (C) Left. Native gel remodeling assay under the conditions of Yan et al. using 50nM SNF2h, saturating ATP, and 15nM cy3-nucleosomes. Remodeling overall is substantially faster likely because of the different conditions used (higher temperature, lower salt concentration, and different components). However, nucleosomes assembled from oxidized H3C110A sCX2 octamers are still remodeled substantially slower than nucleosomes assembled from untreated H3C110A octamers. Nucleosomes assembled from H3C110A sCX2 octamers that were first oxidized and then reduced are remodeled comparably to nucleosomes assembled from untreated H3C110A octamers. Middle. Quantification of the gels on the left along with the indicated observed rate constants. Right. Time-courses shown in the middle panel normalized to the best fit parameters for Y_0_ and Plateau of the exponential decay and zoomed in to the first 10 minutes of the reaction to better evaluate the fits.

**Supplemental Figure 5.**
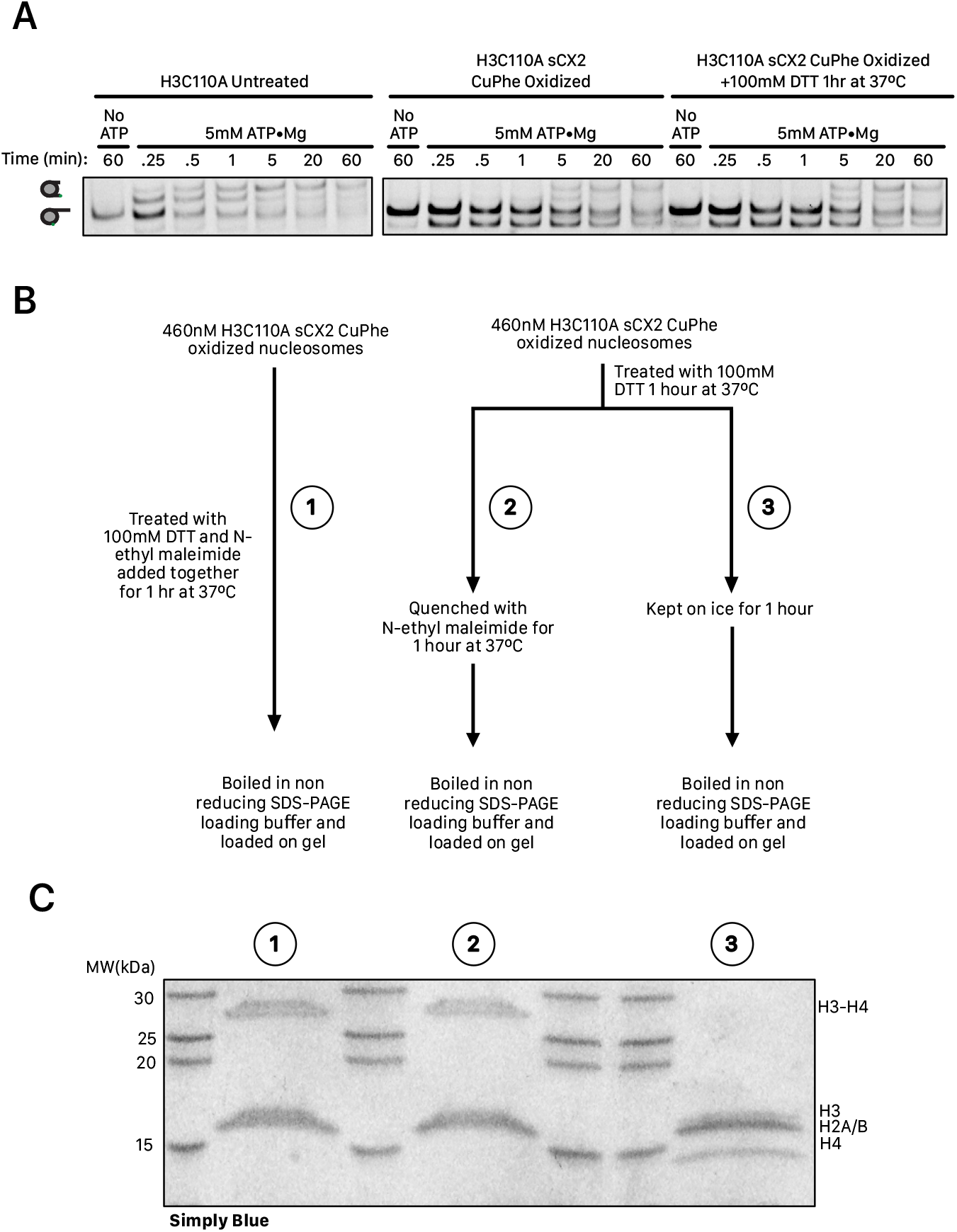
Disulfide reduction is impaired in the context of the nucleosome. (A) Native gel remodeling assay with saturating SNF2h (1μM), saturating ATP, and 15nM cy3-nucleosomes as in Figure 1. Nucleosomes containing the oxidized sCX2 bonds were generated by oxidizing the H3C110A sCX2 octamer using CuPhe, and then assembling nucleosomes. Treatment of these nucleosomes with excess DTT as in Yan et al. fails to reverse the remodeling defect. (B) Scheme for the samples run in C. Nucleosomes treated with DTT were either directly added to non-reducing SDS-PAGE loading buffer or quenched with 500mM N-Ethyl Maleimide freshly dissolved in DMSO (final [DMSO]≈10%(v/v)). Additionally, a condition where N-Ethyl Maleimide and DTT were added simultaneously is included to evaluate the efficacy of the quench. (C) SDS-PAGE of samples treated as in B. Samples with reducing agent quenched prior to running on the gel are near-completely oxidized.

